# Berberine reduce inflammation in RA rats through MCP1/CCR2 pathway

**DOI:** 10.1101/2023.08.09.552722

**Authors:** Wang Ling, Yu Can, Li Meng Ying, Xu Man, Hui Hu

**Affiliations:** Hubei University of Chinese Medicine, Wuhan, Hubei, China; Wuhan Hospital of Traditional Chinese Medicine, Wuhan, Hubei, China

**Keywords:** MCP1/CCR2, Berberine, Rheumatoid Arthritis, chemokine

## Abstract

Rheumatoid arthritis (RA) is a chronic autoimmune disorder characterized by persistent synovitis and systemic inflammation, leading to joint damage and functional disability. Current treatment modalities, although effective, may pose significant side effects. In this study, we investigated the anti-inflammatory properties of berberine, a natural isoquinoline alkaloid, and its potential role in ameliorating RA-associated inflammation in rats through modulation of the monocyte chemoattractant protein-1 (MCP1)/chemokine receptor 2 (CCR2) pathway. A collagen-induced arthritis (CIA) rat model was employed to mimic RA pathophysiology. Rats were treated with berberine or RS504393 for 30 days. Joint swelling, arthritis score, histopathology used to evaluate disease progression and the extent of inflammation. Immunohistochemistry and WB were used to detect the mechanism of action of berberine. Our results demonstrated that berberine treatment significantly reduced joint swelling, and inflammatory factors in CIA rats. Furthermore, berberine downregulated MCP1 and CCR2 expression, implicating the involvement of the MCP1/CCR2 signaling pathway in attenuating RA-associated inflammation. Taken together, our findings suggest that berberine may represent a promising therapeutic candidate for the management of RA and highlight the potential of targeting the MCP1/CCR2 pathway to mitigate inflammation in autoimmune diseases.

## Introduction

Rheumatoid arthritis (RA) is a chronic and systemic autoimmune disease characterized by progressive joint inflammation, stiffness, and pain, ultimately resulting in joint destruction and disability[1]. Epidemiological studies estimate that approximately 1% of the global population is affected by RA, with an increased prevalence in women and the elderly[2]. The etiology of this debilitating disease is multifactorial, involving both genetic and environmental factors[3]. Recent advances in the understanding of RA pathogenesis have highlighted the crucial role of the innate immune system, particularly the involvement of cytokines, chemokines, and other immune mediators in orchestrating inflammatory responses and driving joint destruction[4-6]. While the exact molecular mechanisms underlying RA remain elusive, the interplay between genetic predisposition, aberrant activation of immune cells, and the production of pro-inflammatory factors provide a valuable framework for elucidating potential therapeutic targets.

Chemokines, a family of small, chemoattractant cytokines, play pivotal roles in the initiation and perpetuation of inflammation in RA[7]. These molecular messengers facilitate leukocyte migration to inflammatory sites and contribute to the maintenance of RA by providing a chemotactic gradient for infiltrating immune cells[7]. The dysregulated production and action of chemokines in the synovial tissue contribute to the chronicity of inflammation and ongoing joint destruction in RA[7, 8]. With over 50 members in the chemokine family, each interacts with a specific G-protein-coupled receptor (GPCR) on target cells, making them potential therapeutic targets for RA[9]. A deeper understanding of the chemokine network and its role in RA pathogenesis may uncover novel strategies to modulate the inflammatory process and halt disease progression.

Current treatment options for RA include nonsteroidal anti-inflammatory drugs (NSAIDs), glucocorticoids, disease-modifying antirheumatic drugs (DMARDs), and biological agents[10]. While these therapies alleviate pain and inflammation, they often fail to arrest progressive joint damage or demonstrate long-term efficacy due to their inherent limitations and side effect. Accordingly, there is an ongoing quest for novel and safe therapeutics to target the complex pathology of RA. Traditional Chinese medicine (TCM), with its rich history and broad knowledge base, offers a unique perspective in the identification and development of new RA treatments[11]. Berberine, an alkaloid isolated from several TCM plants, is one such promising candidate, demonstrating potent anti-inflammatory and immunomodulatory properties[12]. Recent studies have revealed that berberine exerts its therapeutic effects in RA through the regulation of multiple molecular pathways, including the inhibition of pro-inflammatory cytokine and chemokine production, suppression of immune cell infiltration, and modulation of intracellular signaling events[12]. Although further research is warranted, berberine represents a promising therapeutic agent in the battle against RA, offering new possibilities for combating this debilitating condition. In our study, we used an experimental RA model and intend to explore the possible molecular mechanism of berberine improvement in RA.

## Method

### Animal and Treatment

A total of 50 male Sprague–Dawley rats, aged 8 weeks and weighing 180-220g, were housed in a 12-hour reversed light-dark cycle with controlled temperature (24 ± 2°C) and humidity (50 ± 5%). Rats had ad libitum access to food and water. CIA was induced in these rats as described previously[13]. CII, emulsified in complete Freund’s adjuvant (1 mg/mL), was intradermally injected at each rat’s tail base at a 0.2 mL volume on day 0 and then boosted with 0.1 mL at day 7. Arthritis symptoms were monitored daily, and paw scores ranked from 0-4. Hind paw volumes were recorded using a plethysmometer. On day 10, rats were allocated into five groups (n=10): Control, Model, RS504393, Berberine, and Berberine+RS504393. Berberine and Berberine+RS504393 groups received daily oral berberine (150 mg/kg). RS504393 and Berberine+RS504393 groups were treated with intra-articular RS504393 (25 μL, 50 ng/mL) using a 27-gauge needle [14], first administered on day 10, then weekly over four weeks. Control and Model groups received a 25 μL intra-articular saline injection.

### Drugs and reagents

Berberine hydrochloride (purity>98%) was purchased from Sigma Aldrich Chemical Co. (St. Louis, USA). Chicken CII and Mycobacterium butyricum were purchased from Sigma-Aldrich (Saint Louis, USA). TNF-α, IL-1β, IL-6, OPG and RANKL ELISA kits were purchased from Meimian (Jiangsu, China). MCP1, IL-6 and IL-8 antibody was purchased from Proteintech (Wuhan, China), CCR2 antibody was purchased from Aviva Systems Biology Co. (San Diego, USA).

### Evaluation of arthritis swelling and arthritis index

Left hind paw swelling in each rat was gauged using a plethysmometer on days 10, 15, 20, 25, 30, 35, and 40 post-FCA immunization. Clinical arthritis severity was evaluated via a scoring system that considered the presence of arthritis in peripheral joints, tail, ears, and both eyes. Cumulative scoring factored in redness and swelling, with scores ranging from 0 to 4: a score of 0 represented no inflammation or swelling, 1 indicated mild erythema or swelling, 2 suggested moderate edema, 3 denoted pronounced edema with limited joint mobility, and 4 signified severe edema accompanied by joint stiffness. The maximum combined arthritis score for each rat’s three paws did not exceed 12 [15].

### Serum IL1-β, TNF-α, IL-6, Receptor activator of nuclear factor-kappa B ligand (RANKL) and osteoprotegerin (OPG) assays

On day 30 post-initial immunization, rats were anesthetized using CO2, 24 hours after the final administration. Blood samples were obtained via the abdominal aorta and left to stand for 30 minutes. Serum was gathered by centrifuging at 1000g for 15 minutes. Serum levels of IL1-β, TNF-α, IL-6, OPG, and RANKL were determined using ELISA kits, following the manufacturer’s protocol. Absorbance was measured at 450 nm using a microplate reader. Concentrations of IL1-β, TNF-α, IL-6, OPG, and RANKL were expressed as picograms or nanograms per milliliter of serum, based on corresponding standard curves.

### Histology, immunohistology, and histological analysis

Rats were euthanized using CO2 and perfused with saline followed by 4% paraformaldehyde. Ankle joint tissues were collected, postfixed for 24 hours in 4% paraformaldehyde, and demineralized in 10% EDTA for six weeks. Specimens were then dehydrated and embedded in paraffin. Longitudinal-oriented sections (4 μm) of the ankle joint were prepared using a paraffin microtome (Finesse 325, Thermo) and processed. For HE staining we used hematoxylin and eosin to stain the sections.

For immunohistochemical staining, primary antibodies used included anti-MCP1 (13.3 μg/mL) and anti-CCR2 (10 μg/mL). Standard immunohistochemical labeling procedures were followed. After dewaxing, antigen retrieval, and blocking with 10% goat serum for 30 minutes at room temperature, sections were incubated with primary antibodies in dilution buffer for 12 hours at 4°C, followed by secondary antibodies (10 μg/mL, Abcam) for 2 hours at room temperature. Nuclei were then stained with hematoxylin and observed under a microscope. Five sections per group were used to observe Trap, MCP1, and CCR2 positive cells in the ankle joint, analyzed using Image J software. For each section, three images were acquired for quantitative analysis. Positive cell percentages were calculated based on total counted synoviocyte and chondrocytes, averaging three fields per slide using Image J. Immunostained images obtained at 400x magnification (n = 5 per group) were employed for histomorphometric analyses. A negative control, with primary antibody replaced by phosphate-buffered saline, was included in each immunostaining experiment.

### Western blot

Rat tissue was lysed in RIPA buffer, and protein concentrations were measured using a BCA protein assay kit (Beyotime, Haimen, China). Protein samples underwent separation on an 8% SDS-PAGE gel before being transferred onto polyvinylidene fluoride (PVDF) membranes. Following a 2-hour blocking with 5% BSA at room temperature, the membranes were incubated overnight at 4°C with primary antibodies: anti-MCP1 (1.33 μg/mL) and anti-CCR2 (1 μg/mL). The membranes were then hybridized with horseradish peroxidase-conjugated secondary antibodies. Upon incubation with enhanced chemiluminescence reagent, protein band images were captured and analyzed using ChemiDoc XRS Image System (Bio-Rad Laboratories). Density values were calculated using Image Lab software, with GADPH as an internal marker; this was expressed as the target protein gray value divided by the overall internal reference gray value.

### Statistical analysis

All data were expressed as mean ± SD and analyzed by GraphPad Prism version 8.0 software. One-way and two-way ANOVA, accompanied by Bonferroni’s post hoc test, were used to assess the differences between groups. A p-value below 0.05 was considered statistically significant.

## Results

### Berberine can alleviate ankle swelling of CIA rat

In order to verify the successful establishment of the model, we firstly observed the gross microscopy of ankle joint and toe of the control and immunized rats and X-ray on the 21st day after the first immunization (Fig.1A). It can be seen that rats in control group did not show toe swelling, and no pathological changes were observed by X-ray. While obvious swelling of ankle and toe could be seen, and the results of X-ray indicated that the hind paws displayed obvious soft tissue swelling, and joint gap stenosis in immunized rats.

**Figure. 1:**
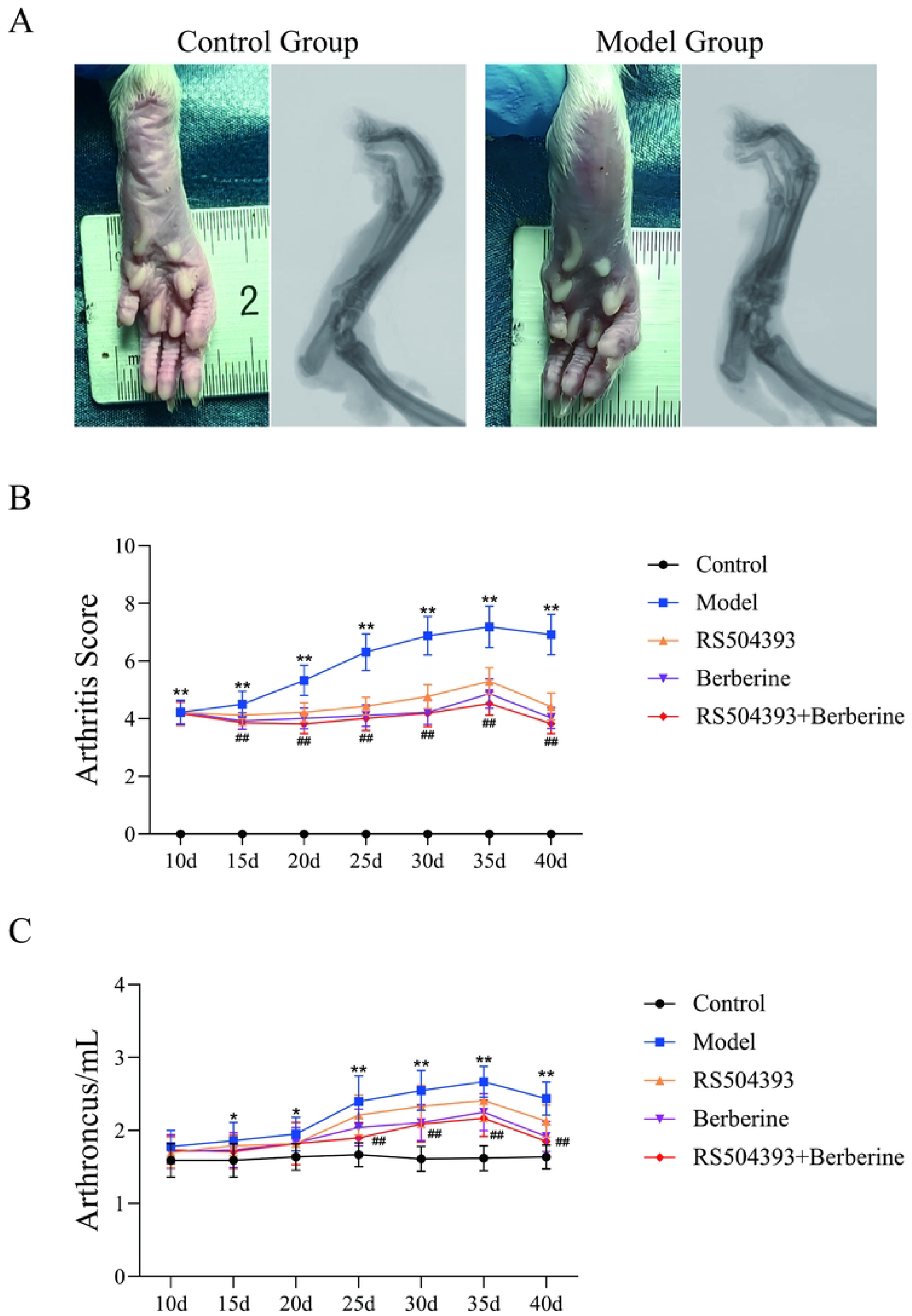
(A) Macroscopic view of ankles and hind paw surface (left), representative images of X-ray radiography of rats in each group (right). (B) Arthritis Score was tested at every 5d since 10^st^ day after immunization (each group: n = 10, two-way ANOVA). (C) Degree of joint swelling was tested by a paw volume plethysmometer at at every 5d since 10^st^ day after immunization (each group: n = 10, two-way ANOVA).

We then detected the hind paw volume by a paw volume plethysmometer and calculate the arthritis index to evaluate severity of arthritis (Figure 1(B) and (C)). In the hind paw volume test, we found that the arthroncus in group model was significantly higher than the control value from the 15d after immunization (P < 0.05). And from the 25d, the arthroncus of the treatment group was significantly lower than that of the model group (P < 0.01). In the arthritis index test, we found that the arthritis score in group model was significantly higher than the control value from the 15d after surgery (P < 0.01). And at the same time, the arthritis score of the treatment group was significantly lower than that of the model group (P < 0.01). But there was no apparent significance between in berberine group and berberine+RS504393 in both the arthritis score and arthroncus.

These results showed that both berberine and RS504393 can protect rats from arthritis after immunization but RS504393 can inhibit the efficacy of berberine.

### Berberine can protect ankle cartilage, reduce osteoclast formation and reduce serum inflammatory factors and RANKL, OPG level in CIA rats

In order to observe the effect of berberine on the pathological morphology of ankle joint in CIA rats, HE and Trap staining was used to evaluated histopathological changes and Osteoclast infiltration (Figure 2(A) and (B)). The trap positive cell was calculated (Figure 2(C)). The results indicated that rats in berberine and RS504393 group showed obvious cartilage protective and Osteoclast formation inhibiting effect. But rats treated with berberin+RS504393 did not show a better effect in cartilage pathological change while compared with rats in berberine model (P > 0.05).

**Figure. 2:**
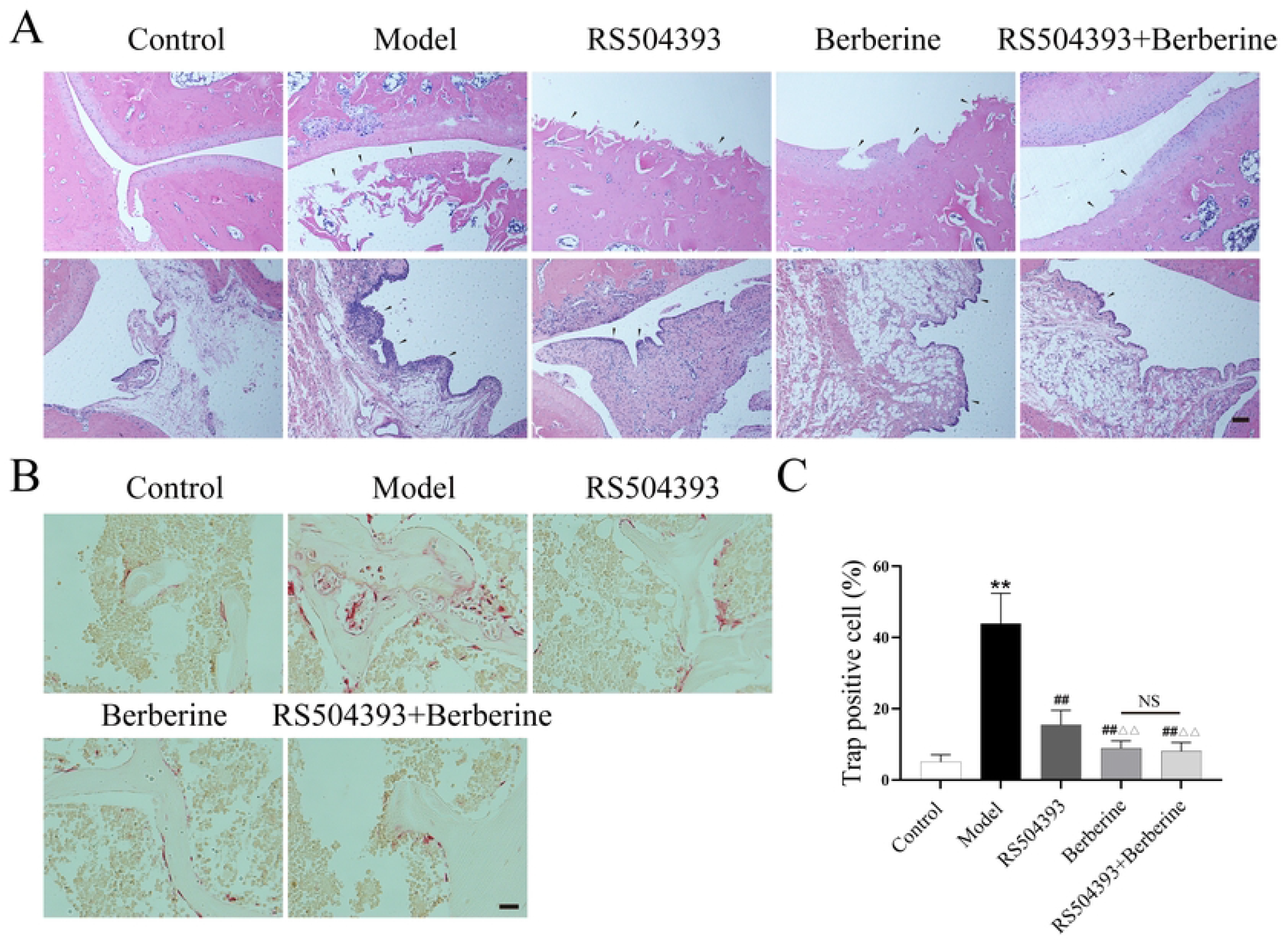
(A) Histological assessment was shown by HE staining (100×, each group: n = 5, Scale bar = 100μm, ANOVA). (B) representative images of Trap staining (400×, each group: n = 5, Scale bar = 20μm, ANOVA). (C) Quantitative analysis of Trap positive cells (each group: n = 5, ANOVA). Values are mean ± SD. ^*^P < 0.05, ^**^P < 0.01: compared with control group. ^#^P < 0.05, ^# #^P < 0.01: compared with model group. ^△^ P < 0.05, ^△△^P < 0.01: compared with RS504393 group. Bonferroni’s post hoc tests were used for multiple comparisons. NS: no significant.

We further detected the level of IL1-β ((Figure 2(A)), TNF-α ((Figure 2(B)), IL-6 ((Figure 2(C)), OPG ((Figure 2(D)), RANKL ((Figure 2(E)) and calculated the ratio of RANKL, OPG ((Figure 2(F)) in serum of rats through ELISA to evaluated the expression level of inflammatory factors and Osteogenic, osteoclast-related factors. The results showed that the expression level of IL1-β, TNF-α, IL-6 and RANKL was greatly elevated in the serum of model rats 40d after immunization while compared with control group (P < 0.01) with OPG reduced apparently (P < 0.01). Berberine and RS504393 can reverse the change above (P < 0.05). Consistent with the results of trap staining, rats treated with berberin+RS504393 did not show a better effect (P > 0.05).

### Berberine reduce the expression of MCP1/CCR2 axis in cartilage and synovia in rats

Chemokines played a vital role in the course of RA. We following observed the expression of MCP1 and CCR2 by immunohistochemical staining (Figure 4(A)) and calculated the MCP1 and CCR2 positive cells ratio in cartilage and synovia (Figure 4(B), (C), (D) and (E)). The results found that rats in model group were significantly higher than control rats in MCP1 and CCR2 positive cell ratio (P < 0.01), and both berberine and RS504393 treatment can reduce the expression of MCP1 and CCR2 (P < 0.01) while compared to rats in model group, but the effects of berberine+RS504393 did not exhibit better efficacy than treated with berberine only (P > 0.05). The results suggested that the effects of berberine in RA are related to CCR2/MCP1 pathway.

**Figure. 3:**
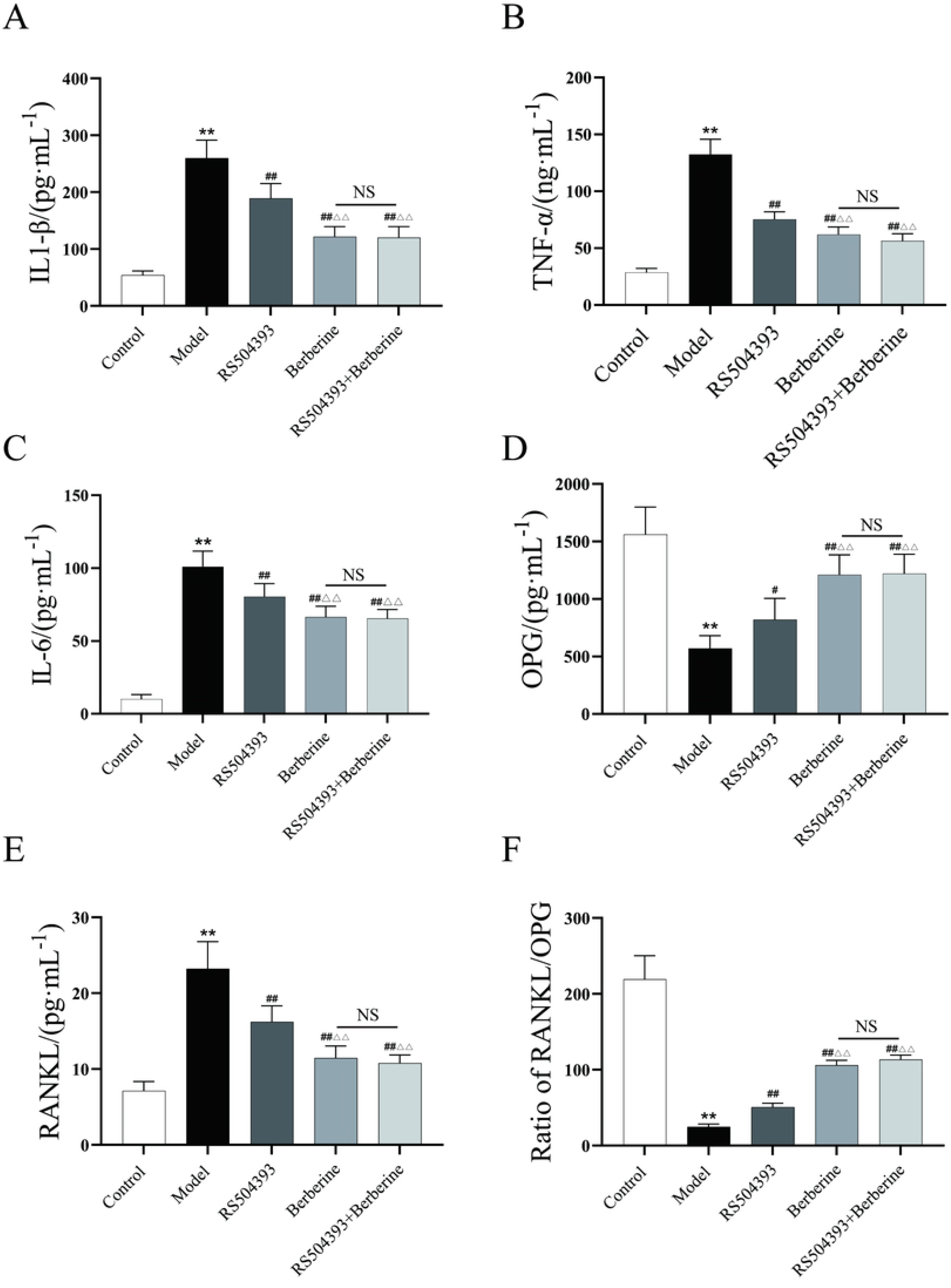
(A, B, C, D, E, F) Quantitative analysis of OPG, IL1-β, TNF-α, IL-6, RANKL, OPG and ratio of RANKL/OPG. (each group: n = 5, ANOVA). Values are mean ± SD. ^*^P < 0.05, ^**^P < 0.01: compared with control group. ^#^P < 0.05, ^##^P < 0.01: compared with model group. ^△^P < 0.05, ^△△^P < 0.01: compared with RS504393 group. Bonferroni’s post hoc tests were used for multiple comparisons. NS: no significant.

**Figure. 4:**
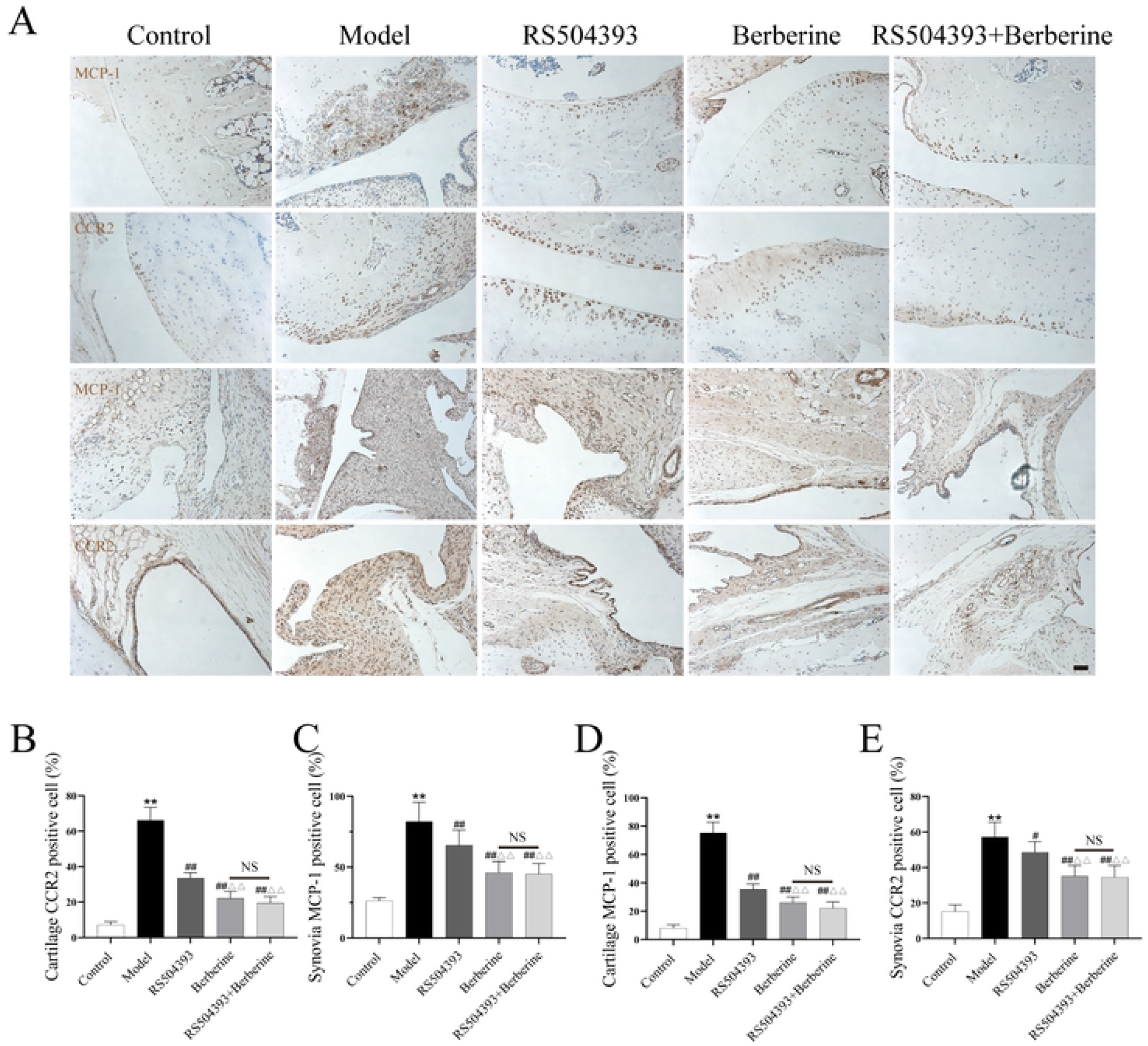
(A) Representative images of immunohistochemistry staining for MCP1 and CCR2 in cartilage and synovia of rats in each group (each group: n = 5, scale bar = 20μm). (B) Quantitative analysis of CCR2 in synovium (each group: n = 5, ANOVA). (C) Quantitative analysis of MCP1 expression level in cartilage (each group: n = 5, ANOVA). (D) Quantitative analysis of MCP1 expression level in synovium (each group: n = 5, ANOVA). Values are mean ± SD. ^*^P < 0.05, ^**^P < 0.01: compared with control group. ^#^P < 0.05, ^##^P < 0.01: compared with model group. ^△^ P < 0.05, ^△ △^ P < 0.01: compared with RS504393 group. Bonferroni’s post hoc tests were used for multiple comparisons. NS: no significant.

### Berberine reduce the expression level of IL6 and IL8 in joint tissue of CIA rat

In RA, local tissue inflammatory factors expression is crucial to the sustained joint inflammation. We analyzed IL6 and IL8 expression by Western blotting (Figure 5(A)) and observed significantly higher levels in the model group than in control rats (Figure 5(B), (C), P < 0.01)). The trend was consistent with the expression of MCP1 and CCR2. Both berberine and RS504393 treatments effectively reduced IL6 and IL8 expression (P < 0.01) compared to the model group. However, combining berberine and RS504393 did not demonstrate increased efficacy over berberine alone (P > 0.05). These findings suggest that berberine mitigates ankle inflammation potentially through the CCR2/MCP1 pathway.

**Figure. 5:**
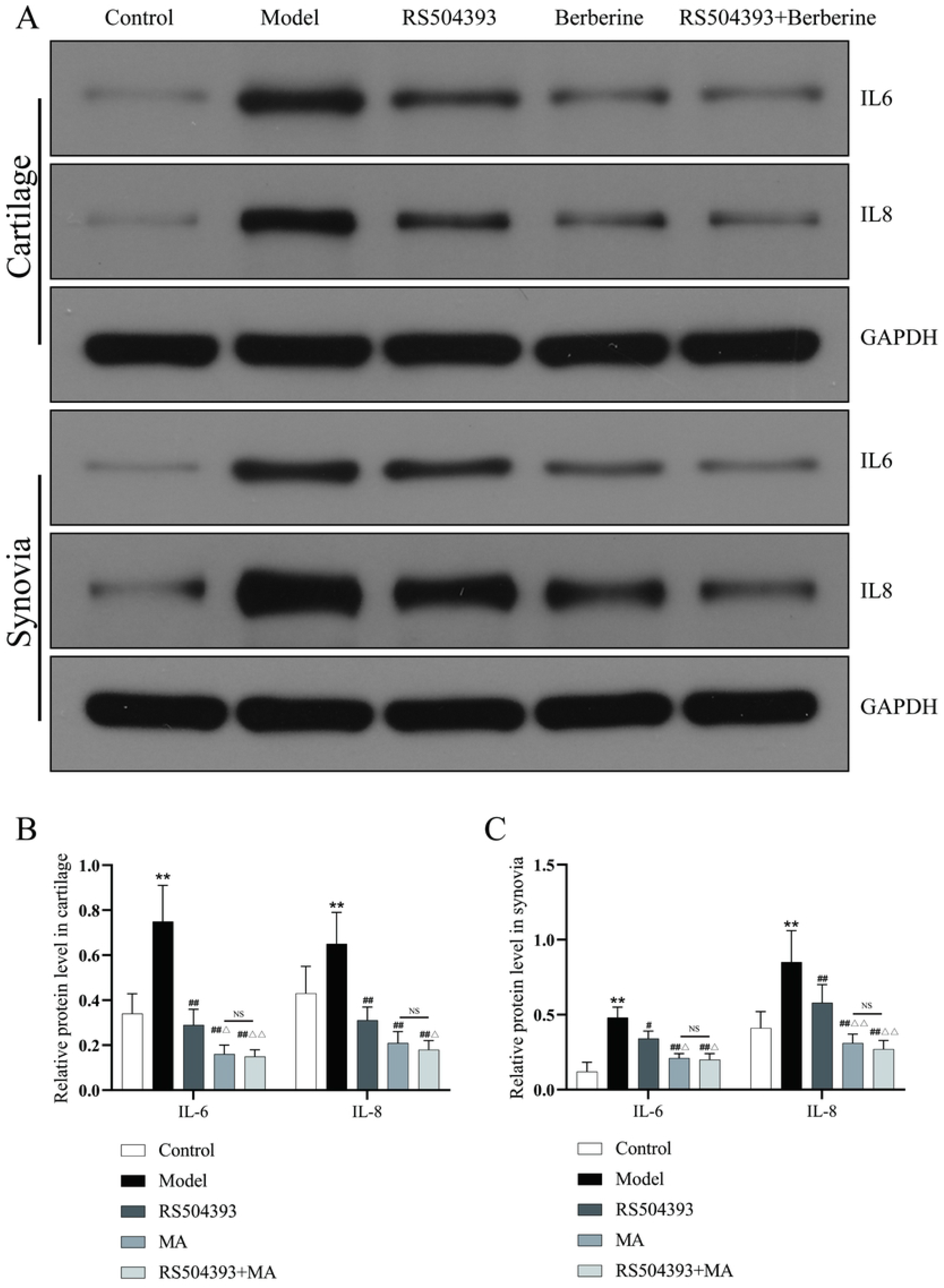
(A) Representative images of western blotting in cartilage and synovia of rats in each group (each group: n = 5). (B) Quantitative analysis of IL-6 and IL-8 in cartilage (each group: n = 5, ANOVA). (C) Quantitative analysis of MCP1 expression level in synovium (each group: n = 5, ANOVA). Values are mean ± SD. ^*^P < 0.05, ^**^P < 0.01: compared with control group. ^#^P < 0.05, ^##^P < 0.01: compared with model group. ^△^ P < 0.05, ^△ △^ P < 0.01: compared with RS504393 group. Bonferroni’s post hoc tests were used for multiple comparisons. NS: no significant.

## Discussion

Rheumatoid arthritis (RA) is a debilitating autoimmune condition characterized by persistent inflammation within the synovial joints, causing cartilage damage and bone erosion[1]. To investigate RA’s underlying biological mechanisms and potential treatments, the collagen-induced arthritis (CIA) rat model has become an invaluable tool[16, 17]. When immunized with either autologous or heterologous type II collagen combined with Freund’s complete adjuvant, rats develop symptoms closely resembling human RA, including joint swelling, erythema, and deformities [18, 19]. The CIA model exhibits many key features consistent with human RA, such as the presence of rheumatoid factors, anti-type II collagen antibodies, and a T helper 1 (Th1) immune response predominance[20, 21]. Moreover, histopathologic findings in this model parallel human RA characteristics, including synovial hyperplasia, pannus formation, and cartilage degradation—further validating its use for studying RA pathogenesis and treatment strategies[22]. Furthermore, the CIA rat model has been employed to evaluate the efficacy of various therapeutic approaches, including disease-modifying antirheumatic drugs (DMARDs), biologic agents, and novel immunomodulators[23, 24]. The progress made using this model has led to a deeper understanding of the cellular and molecular events contributing to RA development and progression, providing essential information for the discovery and development of innovative treatment options. Thus, this model can simulate the pathophysiological process of RA and reinforcing the clinical relevance of our experimental approach.

Berberine is a natural alkaloid present in plants such as Coptis chinensis, Berberis aristata, and Hydrastis canadensis. Over the years, berberine has garnered attention due to its diverse pharmacological effects, including anti-inflammatory, anti-oxidant, and immunomodulatory properties[25]. In recent years, there is growing evidence on the therapeutic potential of berberine for rheumatoid arthritis (RA). Mechanistically, berberine inhibits the nuclear factor-κB (NF-κB) pathway, a crucial signaling cascade involved in the regulation of a variety of pro-inflammatory cytokines, such as tumor necrosis factor (TNF)-α, IL-1β, IL-6, and IL-17[26]. These cytokines play significant roles in RA pathogenesis, and their suppression may limit joint inflammation and damage. Another contributing factor to berberine’s therapeutic effect is its ability to modulate T-cell differentiation, skewing the balance between T-helper 17 (Th17) cells and regulatory T (Treg) cells in favor of Treg cells[27]. This action is particularly relevant to RA, as the imbalance between pro-inflammatory Th17 cells and anti-inflammatory Treg cells has been implicated in the disease’s progression. Furthermore, berberine demonstrates antioxidant effects by increasing the expression of superoxide dismutase (SOD) and catalase (CAT), leading to a reduction in both reactive oxygen species (ROS) production and oxidative stress[28]. Reduced oxidative stress contributes to mitigating inflammation and tissue damage in RA. Berberine also exhibits a suppressive effect on matrix metalloproteinase (MMP) expression[29]. Since MMPs are involved in the degradation of cartilage and other extracellular matrix components in RA, inhibiting their production may have a protective role in joint integrity. Hence, by regulating the NF-κB pathway, modulating T-cell differentiation, reducing oxidative stress, and inhibiting MMP production, berberine could provide a promising therapeutic option for RA management. In this study, we further studied the mechanism of berberine in the treatment of RA, and our results provided a new basis for the clinical treatment of RA-related symptoms.

The monocyte chemoattractant protein-1 (MCP-1) and its receptor, C-C chemokine receptor type 2 (CCR2), play critical roles in the recruitment and activation of monocytes and macrophages in the development of rheumatoid arthritis (RA)[30]. The MCP-1/CCR2 axis has been implicated in the pathogenesis of RA due to its involvement in the recruitment of inflammatory cells to the synovial membrane, leading to joint inflammation and subsequent tissue damage[31]. In RA, elevated MCP-1 levels are observed in synovial fluid and serum, correlating with disease activity[32]. Targeting the MCP-1/CCR2 axis has demonstrated therapeutic potential in animal models of RA, where the inhibition of the MCP-1/CCR2 interaction has been shown to reduce synovial inflammation, suppress joint destruction, and ameliorate clinical symptoms[33]. As a result, the MCP-1/CCR2 axis represents a potential therapeutic target for RA management. In our study, we used RS504393 to intervene on the MCP1/CCR2 axis, and the results showed that blocking the MCP1/CCR2 axis of ankle joint in RA rats had anti-inflammatory effects on RA.

Receptor activator of RANKL and OPG play pivotal roles in bone metabolism, particularly in the context of RA. RANKL is a cytokine that binds to its receptor RANK on osteoclasts, promoting their differentiation and activation, leading to increased bone resorption[34]. OPG, a soluble decoy receptor for RANKL, inhibits RANKL-RANK interaction, thus suppressing osteoclastogenesis and bone resorption[35]. The imbalance between RANKL and OPG has been implicated in the pathogenesis of RA, as excessive bone resorption contributes to joint erosion and deformity[36]. Interestingly, the MCP-1/CCR2 pathway may also modulate the RANKL/OPG axis[37]. In our study, we investigated the effects of berberine and inhibitors on RANKL and OPG, and the results showed that the inhibitory effect of berberine and inhibitors on RANKL/OPG pathway in RA rats may be exerted by inhibiting MCP1/CCR2 pathway. IL-6 and IL-8 are key pro-inflammatory cytokines implicated in the pathogenesis of rheumatoid arthritis (RA). IL-6 contributes to the acute phase response, synovial inflammation, angiogenesis, and joint destruction in RA[38]. IL-8, a chemokine expressed by various cell types in the inflamed synovium, is involved in the recruitment and activation of neutrophils and other immune cells, exacerbating synovial inflammation in RA[39]. The MCP-1/CCR2 pathway also plays a significant role in RA by regulating monocyte and macrophage infiltration into the synovial tissue. Previous studies have suggested that the MCP-1/CCR2 pathway might be interconnected with the IL-6 and IL-8[38, 40]. We detected the expression level of IL-6 and IL-8 in both synovia and cartilage in RA rats and the results were consistent with those of chemokines.

In conclusion, our study provides evidence that berberine exhibits significant anti-inflammatory effects in CIA rat models by modulating the MCP1/CCR2 signaling pathway. This novel finding presents berberine as a promising therapeutic agent for managing RA, potentially reducing its progression and improving the quality of life for affected patients. Further research is warranted to elucidate the precise molecular mechanisms underlying berberine’s anti-inflammatory properties and to assess its efficacy and safety profile in human subjects with RA, thereby paving the way for its potential integration into the existing repertoire of RA treatments.

